# TMEM170 family proteins are lipid scramblases that physically associate with bridge lipid transporters BLTP1/Csf1

**DOI:** 10.1101/2025.04.30.651444

**Authors:** Cristian Rocha-Roa, Gurpreet Sidhu, Paige Chandran Blair, Daniel Álvarez, Michael Davey, Elizabeth Conibear, Stefano Vanni

## Abstract

Bulk lipid transport between organelles has been proposed to involve the partnership between bridge lipid transport proteins and membrane-embedded lipid scramblases. However, for almost all BLTPs, such physical association has not been fully described, and, in most cases, the identity of the scramblases is unknown. Here, we identify TMEM170 family proteins as endoplasmic reticulum lipid scramblases that physically interact with BLTP1/Csf1 proteins. This finding opens new avenues to understand the complex mechanism involved in lipid transport at membrane contact sites.

## Main

Several bridge-like lipid transport proteins (BLTPs) of the Vps13/Atg2 family have been shown to associate with integral membrane proteins possessing lipid scrambling activity. Examples include Atg2-Atg9^1,2^ at the autophagosome membrane, ATG2-TMEM41B/VMP1^3,4^ at the endoplasmic reticulum (ER) membrane, Vps13-Mcp1 at the vacuole-mitochondria membrane contact site (MCS)^5^, VPS13A and Xk at the plasma membrane^6,7^, and, very recently, Vps13-Any1 at the ER-Early endosome MCS^8^. BLTPs are proposed to cooperate with lipid scramblases in both donor and acceptor membranes to facilitate lipid uptake and release, and to promote membrane growth by breaking down the intrinsic leaflet asymmetry of lipid transport, since, otherwise, lipid would be transported only between the donor and acceptor cytosolic leaflets. However, for the vast majority of known BLTPs, the corresponding scramblases remain unknown and uncharacterized, and the generality of such a molecular arrangement requires further validation. Additionally, although the close relationship between a BLTP and a scramblase has been proposed to be critical for bulk lipid transport, the lack of well-characterized 3D structures detailing the physical interaction between these two components makes it hard to address the interplay between them and to evaluate a potential cooperative mechanism.

To identify new lipid scramblases that associate with BLTPs, we first noticed that in a previous screen for yeast growth under high hydrostatic pressure and/or low temperature^9,10^, perturbations that increase lipid order and decrease membrane fluidity, the lack of ER protein Ypr153w/May24 phenocopied the loss of either the BLTP Csf1 (BLTP1 in human) or Ymr126c/Dlt1^9^. Notably, the Dlt1 homologue in *C. elegans*, Spigot, was recently shown to physically associate with the *C. elegans* homologue of BLTP1 and Csf1, LPD-3^11^. These data point to the ER-resident protein Ypr153w/May24 as a potential partner of the BLTP Csf1.

Bioinformatics analyses indicate that Ypr153w/May24 is the homologue of human TMEM170A/B. They are relatively small proteins (∼15.5 kDa) consisting exclusively of a core transmembrane (TM) helical bundle composed of three TM helices (Fig. 1a). Additionally, human TMEM170A has been proposed as a regulator of ER and nuclear envelope morphogenesis via a yet-uncharacterized mechanism^12^, suggesting a possible link with lipid homeostasis.

**Figure 1.**
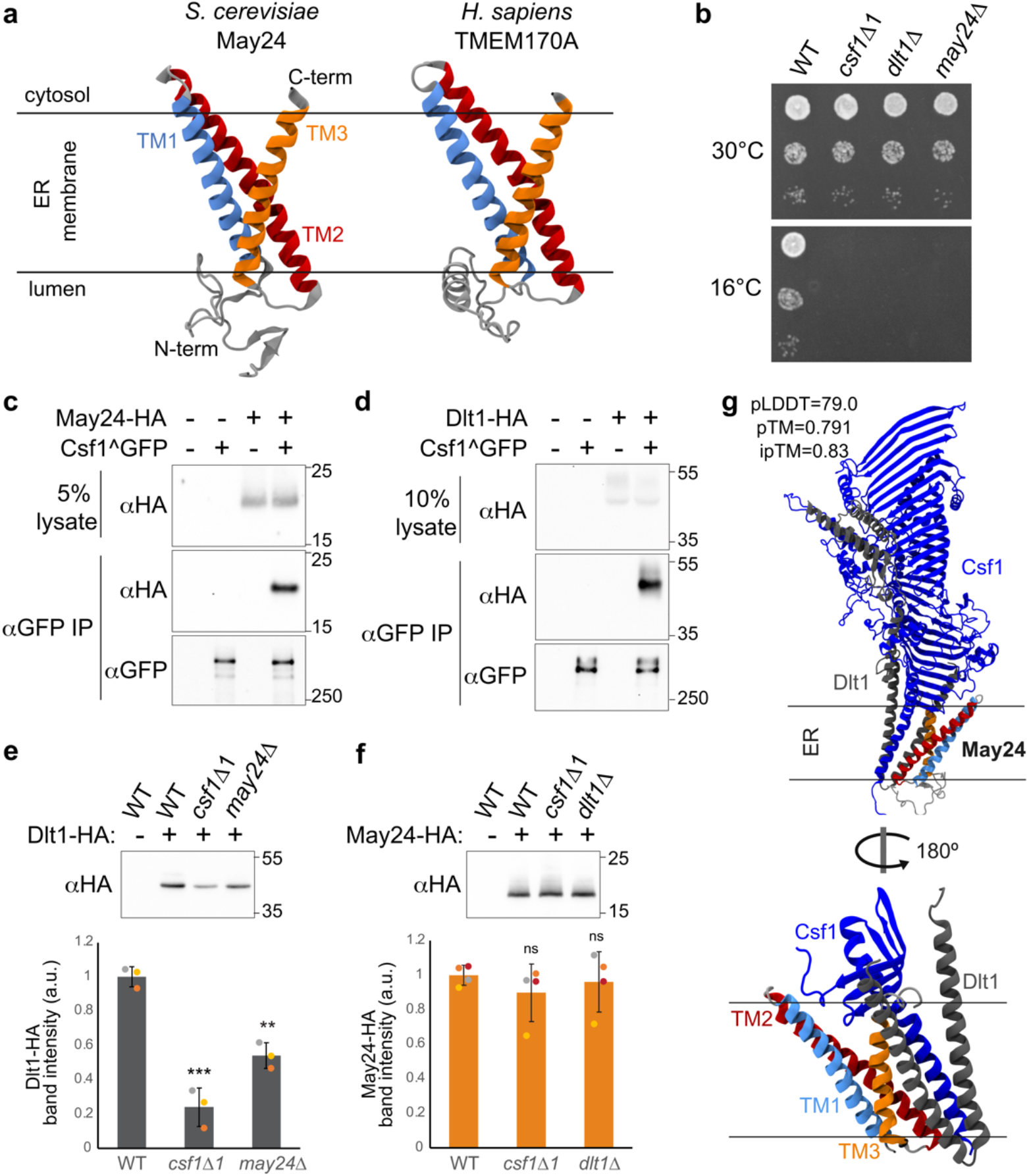
**a.** AF structural models for full-length yeast ER protein May24 (left panel) and its human homologue TMEM170A (right panel). The three TM helices are shown in light blue, red and orange, respectively. In the yeast model structure, the N-term and C-term are labeled as visual references. **b**. Mutants lacking May24 have a cold sensitive phenotype comparable to that of *csf1Δ1* and *dlt1Δ* mutants. **c**. May24 co-purifies with Csf1 from cell lysates. Both proteins are epitope-tagged at their endogenous chromosomal locations. Molecular weight (kDa) is indicated on the right. **d**. Immunoprecipitation of Csf1 efficiently copurifies Dlt1 from cell lysates. **e**. Dlt1 is destabilized in the absence of Csf1 or May24. Western blot analysis of endogenously tagged Dlt1-HA, n=3, normalized to the average value in the wild type (WT) strain. One-way ANOVA with Tukey’s multiple comparison test; *** : p=0.000084; ** : p=0.0014. Error bars indicate standard deviation. **f**. The stability of May24-HA is unaffected by loss of Csf1 or Dlt1. Western blot analysis of endogenously tagged May24-HA, n=4, normalized to the average value in the wild type (WT) strain. One-way ANOVA; ns: p=0.62. Error bars indicate standard deviation. **g**. Upper panel: AF model of the yeast complex formed by Csf1 (blue, sequence from Met1 to Ala1200), Dlt1 (gray, sequence from Met1 to Arg250) and May24 (same color code as in **a**). Lower panel: close-up view of the TM part of the complex.

To better characterize the role and mechanism of TMEM170 family proteins (Pfam: PF10190) in BLTP-mediated lipid transport, we used genetic, biochemical and computational approaches. First, we confirmed that deletion of *MAY24, CSF1* or *DLT1* in yeast resulted in a similar degree of cold sensitivity (Fig. 1b), suggesting they have closely related functions. We next tested whether the yeast TMEM170 protein May24 physically associates with the BLTP Csf1. Immunoprecipitation of Csf1 efficiently co-purified both May24 (Fig. 1c) and Dlt1 (Fig. 1d). The steady-state level of Dlt1 was also reduced in the absence of either Csf1 or May24 (Fig. 1e), consistent with the model that Csf1, Dlt1 and May24 form a trimeric complex. Because the stability of May24 was unaffected by the loss of Csf1 or Dlt1 (Fig. 1f), this protein may also exist in a separate pool.

AlphaFold (AF) modeling of the complex formed by Csf1 together with Dlt1 and May24 places the TMEM170 family protein May24 in direct contact with the TM domain of Csf1 (Fig. 1g). A similar predicted interaction pattern is observed for the corresponding human complex, comprised of BLTP1, TMEM170A and C1orf43, the human homologue of yeast Dlt1 (Fig. S1a and Table S3). Notably, this arrangement is consistent with the cryo-EM density observed in a recent work from Kang and co-workers^11^ where the structure of the N-terminus of *C. elegans* LPD-3 was solved using single-particle cryo-EM at high resolution (2.7 Å). There, in addition to the integral membrane density arising from the sole TM helix of LPD-3, four additional TM helices could be observed, with one of these helices assigned to Spigot, the homologue of yeast Dlt1 and human C1orf43. Our AF models of the complexes (Fig. 1g, Fig. S1 and Table S3) place the three TM helices of the TMEM170 protein exactly at the spot where the additional unassigned membrane density is observed in the cryo-EM map, consistent with our biochemical analyses. As a control, AF predictions for other BLTPs similar to Csf1 in *S. cerevisiae*, Hob1 (UniProt ID: Q06179) and Hob2 (UniProt ID: Q06116), did not show interaction between the May24/Dlt1 dimer and the TM helix of neither Hob1 nor Hob2.

To further support our biochemical and bioinformatics observations, we next processed the single-particle cryo-EM map of the *C. elegans* LPD-3 complex (EMDB ID: 45399) (Fig. 2a, left panel) using ModelAngelo, a machine-learning approach for automated atomic model building in cryo-EM maps^13^. In addition to the homologues of yeast Csf1 and Dlt1 (LPD-3 and Spigot in *C. elegans*, respectively), ModelAngelo independently identified the *C. elegans* homologue of May24/TMEM170A, CELE_F43G9.13 (Uniprot ID: Q95QG3, which we will refer to here as Tmem-170), in the additional transmembrane density (Fig. 2a, center panel). Comparison between the ModelAngelo-predicted structure and the AF model (Fig. 2a, right panel) further supports our observation of TMEM170 proteins as direct transmembrane partners of BLTP1/Csf1 proteins (Fig. 2a and Table S3).

**Figure 2.**
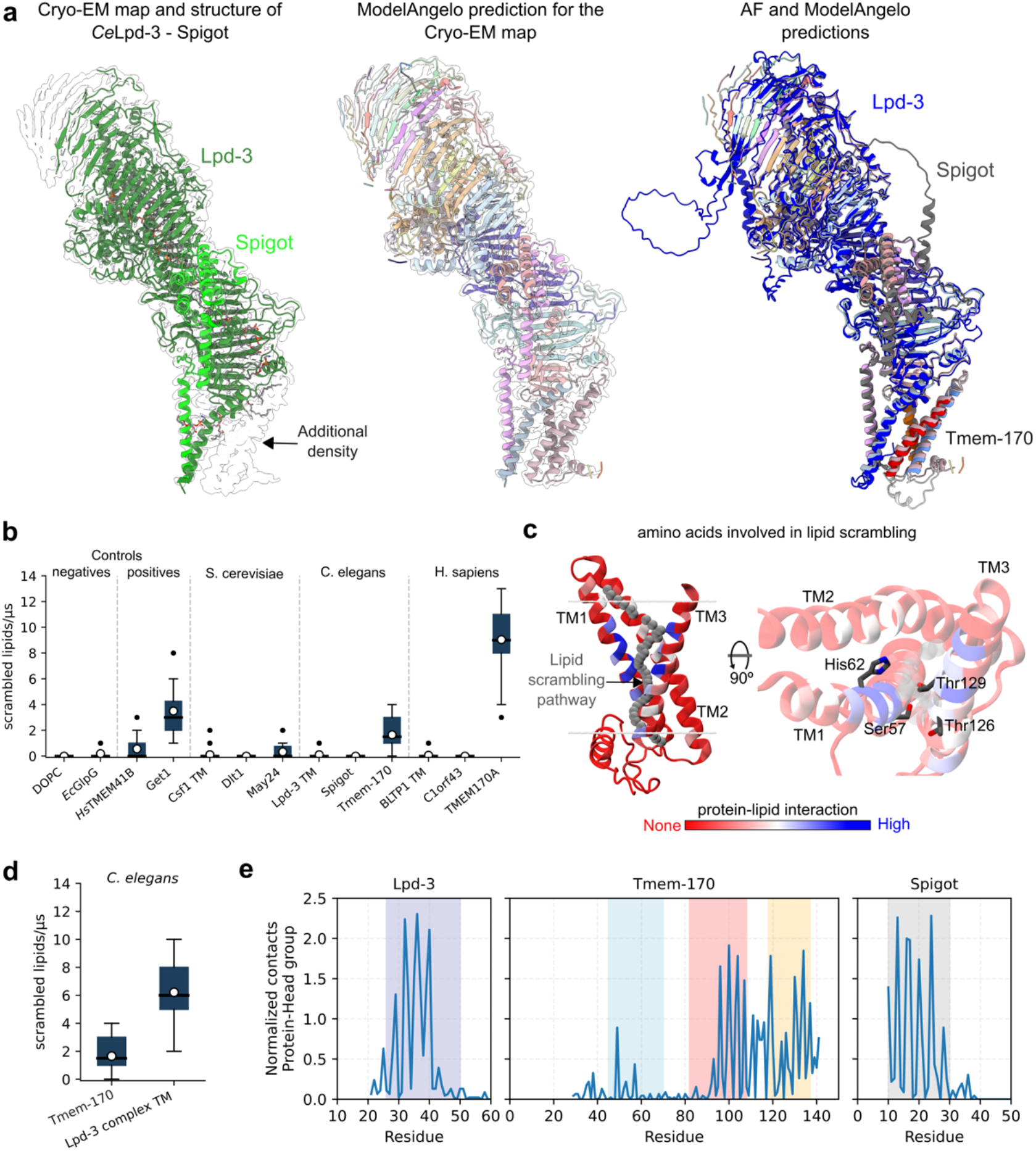
**a.** Computational-assisted identification of *C. elegans* Tmem-170 protein as part of a heterotrimeric complex involved in bulk lipid transport. Left panel: CryoEM map and corresponding atomistic structure of the complex formed by Lpd-3 (green), Spigot (light green), and an unresolved unknown protein from reference ^11^. Center panel: ModelAngelo output after processing the Cryo-EM map in the left panel. Right panel: structural alignment of the AF prediction for the *C. elegans* complex and the ModelAngelo prediction. The AF prediction is colored as in Figure 1g. **b**. *In silico* lipid scrambling assay for the yeast (Csf1, Dlt1 and May24), worm (Lpd-3, Spigot, and Tmem-170) and human (Bltp1, C1orf43 and TMEM170A) proteins. Data for positive (*Hs*TMEM41B and *Sc*Get1) and negative (DOPC and *Ec*GlpG) controls are taken from reference^14^. For all systems, a total of 46 data points were used to build each boxplot. The blue boxes represent the interquartile region. The solid black line inside each box represents the median of the data while the whiskers represent the data range. Averages are shown as white circles. Outliers are shown as black dots. **c**. Main amino acids involved in lipid scrambling for the human TMEM170A protein. Left panel: interactions between the protein and the scrambled lipids are shown in a color scale from blue (high frequency of interactions) to red (no interactions). The averaged lipid scrambling pathway is presented as a continuous stalk of gray beads. Right panel: the main polar amino acids involved in lipid scrambling are labeled and shown as sticks. The transmembrane region and labels for each TM helix are shown as visual reference. **d**. *In silico* lipid scrambling assay for the worm Tmem-170 alone (left) and for worm Tmem-170 in complex with the TM domains of Lpd-3 and Spigot (right). Statistical analyses as in panel **(b). e**. Interaction between the head groups of the scrambled lipids and each component of the heterotrimeric complex from *C. elegans* (Left panel: Lpd-3; Center panel: Tmem-170; Right panel: Spigot). The TM segments of Lpd-3 and Spigot are highlighted with blue and gray shadows, respectively; while the three TM segments of Tmem-170 are highlighted as in panel **(a)** and Figure **1a**.

Next, we used a well-established *in silico* assay based on molecular dynamics (MD) simulations^14^ to assess lipid scrambling activity by TMEM170 family proteins. Our results indicate that while neither BLTPs (Csf1, Lpd-3, BLTP1) nor Dlt1/Spigot/C1orf43 alone can scramble lipids, TMEM170 family proteins can scramble lipids with similar rates as other ER scramblases tested with both *in silico* and *in vitro* assays (Fig. 2b). Specifically, May24 has comparable activity with the bona fide human ER scramblase TMEM41B^3,14^, while human TMEM170A and worm Tmem-170 have scrambling activity comparable with that of the yeast ER scramblase Get1^14^. To further generalize our observation, we built a phylogenetic tree for TMEM170 family proteins using available reviewed sequences in Swiss-Prot (Fig. S2a) and assessed *in silico* lipid scrambling for all the TMEM170 proteins we identified. These data confirm that TMEM170 family proteins can scramble lipids and show that lipid scrambling appears more pronounced in mammals TMEM170A proteins (Fig. S2b). Additionally, our AF predictions suggest that all simulated TMEM170 proteins may form similar heterotrimeric complexes, with high confidence scores. This could indicate that a conserved protein machinery for bulk lipid transport exists across cells of varying complexity (Table S3).

Analysis of the lipid scrambling pathway in the monomeric structure of human TMEM170A (Fig. 2c) indicates that scrambling is promoted by the presence of multiple polar residues inside the TM region protein (Ser57, His62, Thr123, Thr126, and Thr129). Comparison between the monomeric structure and the full trimeric complex in *C. elegans*, however, reveals that while scrambling activity can be observed in both systems (Fig. 2d), in the trimeric complex all proteins contribute to the scrambling pathway (Fig. 2e and Fig. S3), suggesting a potential cooperative mechanism linking lipid scrambling with transport by the BLTP.

Taken together, our data indicate that TMEM170 family proteins are lipid scramblases that physically associate in the ER with BLTPs Csf1/BLTP1/Lpd-3. Our data support the current model of tight interplay between bridge lipid transporters and lipid scramblases, and pave the way for two lines of investigations: the further characterization of the lipid transport mechanism by lipid scramblase-associated BLTPs, and the identification of additional functions beyond their association with BLTPs for lipid scramblases of the TMEM170 family.

## Acknowledgments

We thank K. Reinisch and W. Kukulski for useful discussions and comments regarding this manuscript, and to S. Clark for sharing the cryo-EM map of the *C. elegans* Lpd-3 complex before public release. We gratefully acknowledge support from the Swiss National Science Foundation (grants CRSII5_189996 to SV), the European Research Council under the European Union’s Horizon 2020 research and innovation program (grant agreement no. 803952, to SV), the Canadian Institutes of Health Research (grants OGB-177941 and PJT-180544 to EC), and a BC Children’s Hospital Research Masters studentship (to PCB). This work was supported by grant s1269 from the Swiss National Supercomputing Centre and by funding from the Canada Foundation for Innovation (Leading Edge Fund 30636).

## Data Availability Statement

The cryo-EM map of the *C. elegans* complex formed by Lpd-3, Tmem170 and Spigot can be found at the Electron Microscopy Data Bank (EMDB) under accession code 45399. All MD trajectories can be found in Zenodo, at https://doi.org/10.5281/zenodo.15272488. The in-house script used to carry out the lipid scrambling analysis, as well as data for the negative (pure DOPC membrane and *Ec*GlpG) and positive (*Sc*Get1 and *Hs*TMEM41B) controls can be found at https://doi.org/10.5281/zenodo.10475371.

## Methods

### 3D structure predictions

Identification of protein homologues was made through multiple sequence alignments carried out by the HHpred web server^15^ and the BLAST tool available in UniProt^16^. The sequences of the identified proteins were retrieved from UniProt (Last consulted on February 24, 2025) and subsequently used for structure prediction with AlphaFold2^17,18^ via ColabFold^19^. For this, 12 recycles and 3 different seeds were used.

A 3D structure for the *C. elegans* complex was generated by processing the Cryo-EM density map (EMDB ID: 45399) using ModelAngelo^13^. Due to the high resolution of the Cryo-EM map (2.7 Å) the option “*build_no_seq”* option was used (no sequences were used as input).

Structural alignments were carried out with the *MatchMaker* tool of the software UCSF Chimera 1.16 and the Needleman-Wunsch alignment algorithm.

The transmembrane orientation for the predicted structure of the single proteins and complexes were locally computed with PPM^20^.

### In silico scrambling assay

Lipid scrambling by TMEM170 proteins was tested using coarse-grained molecular dynamics (CGMD) simulations. All simulations were performed using the software GROMACS^21^ with the CG Martini 3 force field^22^. To run CG simulations, the all-atom structures predicted for all the proteins and complexes were converted into CG resolution using the *Martinize2* script^23^. To preserve the 3D structure of the proteins, an elastic network was employed using a force constant of 500 kJ mol^−1^ nm^−2^ and an elastic bond upper cut-off of 0.9 nm. In the cases of more than one protein chain the elastic networks were merged. Using the *Insane* script^24^, the CG proteins were embedded into a membrane composed of pure DOPC phospholipids and then solvated with 0.15 M NaCl solution plus additional Na^+^ or Cl^−^ions to neutralize protein charges.

Equilibration consisted of two energy minimization stages using the steepest-descent algorithm, for a total of 25000 steps or until the maximum force was smaller than 10 kJ mol^−1^ nm^−1^; positional restraints were applied on the backbone beads of the proteins with a force constant of 1000 kJ mol^−1^ nm^−2^. Next, two 1-ns long NVT simulations with a time- step of 2 fs were carried out. For the first NVT equilibration run, positional restraints of 1000 kJ mol^−1^ nm^−2^ and 200 kJ mol^−1^ nm^−2^ were used on the protein backbone beads and phosphates, respectively, and of 900 kJ mol^−1^ nm^−2^ and 190 kJ mol^−1^ nm^−2^ in the second run. Then, five NPT equilibrations were carried out by slowly increasing the time step and decreasing positional restraints.

Finally, a production run of 25 µs using a time step of 20 fs was carried out. The temperature was defined at 310 K and controlled with the v-rescale thermostat^25^ with a coupling constant of 1 ps. Protein, lipids and solvent were coupled separately. The pressure was defined at 1 bar and was controlled with a semi-isotropic Parrinello-Rahman barostat^26^ and a coupling constant of 12 ps. Electrostatic interactions were computed using the reaction-field approach with a cut-off of 1.1 nm. Van der Waals interactions were computed using the cut-off scheme with a cut-off distance of 1.1 nm. The Potential-shift-verlet was used with a Verlet cut-off scheme for both types of interactions. The Verlet list scheme was used to update the particle neighbor list every 20 steps. Two independent replicas (different initial velocities) were performed for each system.

Lipid scrambling was quantified by tracking the tilt angles (*θ*) of each lipid with respect to the membrane plane normal throughout the simulation. A flip (movement from the upper to the lower leaflet) or flop (movement from the lower to the upper leaflet) event was considered when the lipid exhibited a change to tilt angles *θ* ≥ 130° or *θ* ≤ 50°, respectively. The angles were computed every nanosecond with the *gmx_gangle* tool, and events were reported as events/μs, omitting the first 2μs for equilibration. A more detailed and complete description of the analysis can be found in the original study^14^. The in-house script used to carry out the analysis and the data for the negative (pure DOPC membrane and *Ec*GlpG) and positive (*Sc*Get1 and *Hs*TMEM41B) controls can be found in https://doi.org/10.5281/zenodo.10475371.

### Phylogenetic tree analysis

Protein sequences of the TMEM170 proteins were retrieved from Uniprot, and only the Reviewed sequences were selected (Swiss-Prot). By using the Clustal Omega web server (https://www.ebi.ac.uk/jdispatcher/msa/clustalo), the sequences were aligned, and the corresponding phylogenetic tree was built.

### Yeast Strains and Plasmids

Yeast strains, plasmids and primers used in this study are described in Supplemental Table S1. Plasmids were made by Goldengate assembly in *Escherichia coli* and fully sequenced. Homologous recombination-based integration of PCR products^27,28^ was used for gene deletions and tagging with the 3xHA epitope, which were confirmed by PCR of genomic DNA. CRISPR-Cas9-based genome editing approaches were used to tag endogenous Csf1 at an internal site (residue 804) with the Envy variant of GFP^29^. A custom linearized Cas9-gRNA plasmid targeting *CSF1* sequences was co-transformed with PCR-amplified donor DNA. CRISPR-Cas9 edited strains were validated by DNA sequencing and grown without selection to lose gRNA plasmids and restore auxotrophy. To create a *csf1* loss of function mutation that does not perturb expression of the neighboring *GAA1* gene (*csf1Δ1*), codons for residues 1003-2018 were replaced by a NATMX4 selectable marker. All strains and plasmids are available on request.

### Western blot and co-immunoprecipitation

For western analysis of protein stability, yeast were grown overnight at 30°C in complete yeast peptone dextrose media (YPD) (1% [w/v] yeast extract, 2% [w/v] Bacto-Peptone, 2% [w/v] glucose, 0.032% [w/v] tryptophan), then diluted and grown for 4 hours to logarithmic phase. 10 OD_600_/mL equivalent of cells were harvested, placed at -80 °C for 5 min, and resuspended in 100μL Thorner buffer (40mM Tris-Cl pH 6.8, 8M urea, 5% SDS, 1% β-mercaptoethanol and 0.4 mg/mL bromophenol blue). Cells were lysed by vortexing with an equal volume of 0.5 mm diameter glass beads. Proteins were denatured at 30°C for 5 min and separated on 10% or 13% SDS-PAGE gels.

For co-immunoprecipitation experiments, 25 OD_600_/mL equivalent of cells were harvested, converted to spheroplasts by digesting cell walls with Zymolyase (SK1204911; MJS BioLynx, Brockville, Canada) and stored at -80 °C. Spheroplasts were thawed in 500μl lysis buffer (50 mM HEPES pH 7.4, 1% Nonaethylene glycol monododecyl ether [C12E9], 50 mM NaCl, 1 mM EDTA, 1 mM PMSF, and 1X yeast/fungal Protease Arrest). 50μl of lysate was set aside and Laemmli Sample Buffer (50 mM Tris-Cl pH 6.8, 2% SDS, 10% glycerol, 0.001% bromophenol blue, 2% β-mercaptoethanol) was added from a 5x stock solution. The remaining lysate was incubated with polyclonal rabbit anti-GFP (EU2; Eusera) for 1 h followed by Protein A-Sepharose beads (GE Healthcare) for 1 h at 4°C. Washed beads were resuspended in 50μl of 2X Laemmli Sample Buffer. For the May24/Csf1 coIP, proteins were eluted by heating at 70°C for 5 min. For the Csf1/Dlt1 coIP, samples were first heated to 37°C and loaded on a gel to detect Dlt1-HA, and subsequently heated to 70°C before loading on a second gel to detect Csf1^GFP.

After separation on SDS-PAGE gels, proteins were transferred to nitrocellulose membranes (50-206-3328; Fisher Scientific) and blotted with either mouse monoclonal α-GFP (clones 7.1 and 13.1, 11814460001; Roche; RRID:AB_390913), HA.11 (clone 16B12, MMS-101R; Covance; RRID:AB_291262) or α-PGK1 (clone 22C5D8; Thermo Fisher Scientific; RRID:AB_2532235) antibodies, followed by horseradish peroxidase-conjugated polyclonal goat α-mouse (115–035-146; Jackson ImmunoResearch Laboratories; RRID: AB_2307392) or horseradish peroxidase-conjugated polyclonal goat anti-rabbit (170-6515; Biorad; RRID: AB_11125142) antibodies. Blots were developed with ECL chemiluminescent reagents (RPN2106, Cytiva) and imaged on an iBright (Invitrogen) imaging system. Densitometry of scanned films was performed in Fiji. Replicates represent independent analyses of the indicated strains. Statistical analysis was carried out in Microsoft Excel version 16.95.4 using the XRealStats-Mac package. Data distribution was assumed to be normal, but not formally tested.

### Growth assays

For plate-based growth assays, overnight cultures were diluted and grown 4 hours to a concentration of 1-2 OD^600^/mL. 10-fold serial dilutions were spotted onto two sets of YPD plates that were incubated at either 16°C for 4 days, or 30°C for 24 h.

## Supplementary information

**Table S1.** Yeast strains used in this study.

| Strain Name | Genotype | Source |
| --- | --- | --- |
| BY4741 | *MATa his3Δ1 leu2Δ0 met15Δ0 ura3Δ0* | Open Biosystems |
| GSY1 | BY4741 *MAY24-3HA::HIS3MX6* | This study |
| GSY2 | BY4741 *dlt1Δ::KANMX6 MAY24-3HA::HIS3MX6* | This study |
| GSY3 | BY4741 *csf1Δ1(1003::NATMX4::2108) MAY24-3HA::HIS3MX6* | This study |
| GSY7 | BY4741 *DLT1-3HA::HIS3MX6* | This study |
| GSY8 | BY4741 *may24Δ::KANMX6 DLT1-3HA::HIS3MX6* | This study |
| GSY9 | BY4741 *csf1Δ1(1003::NATMX4::2108) DLT1-3HA::HIS3MX6* | This study |
| MDY1340 | BY4741 *CSF1(1-804)-GFP<sup>Envy</sup>-CSF1(805-end)* | This study |
| GSY13 | BY4741 *CSF1(1-804)-GFP<sup>Envy</sup>-CSF1(805-end) DLT1-3HA::HIS3MX6* | This study |
| GSY15 | BY4741 *CSF1(1-804)-GFP<sup>Envy</sup>-CSF1(805-end) MAY24-3HA::KANMX6* | This study |
| SSY864 | BY4741 *csf1Δ1(1003::NATMX4::2108)* | This study |
| GSY33 | BY4741 *dlt1Δ::KANMX6* | This study |
| SSY872 | BY4741 *may24Δ::KANMX6* | Open Biosystems |

**Table S2.**
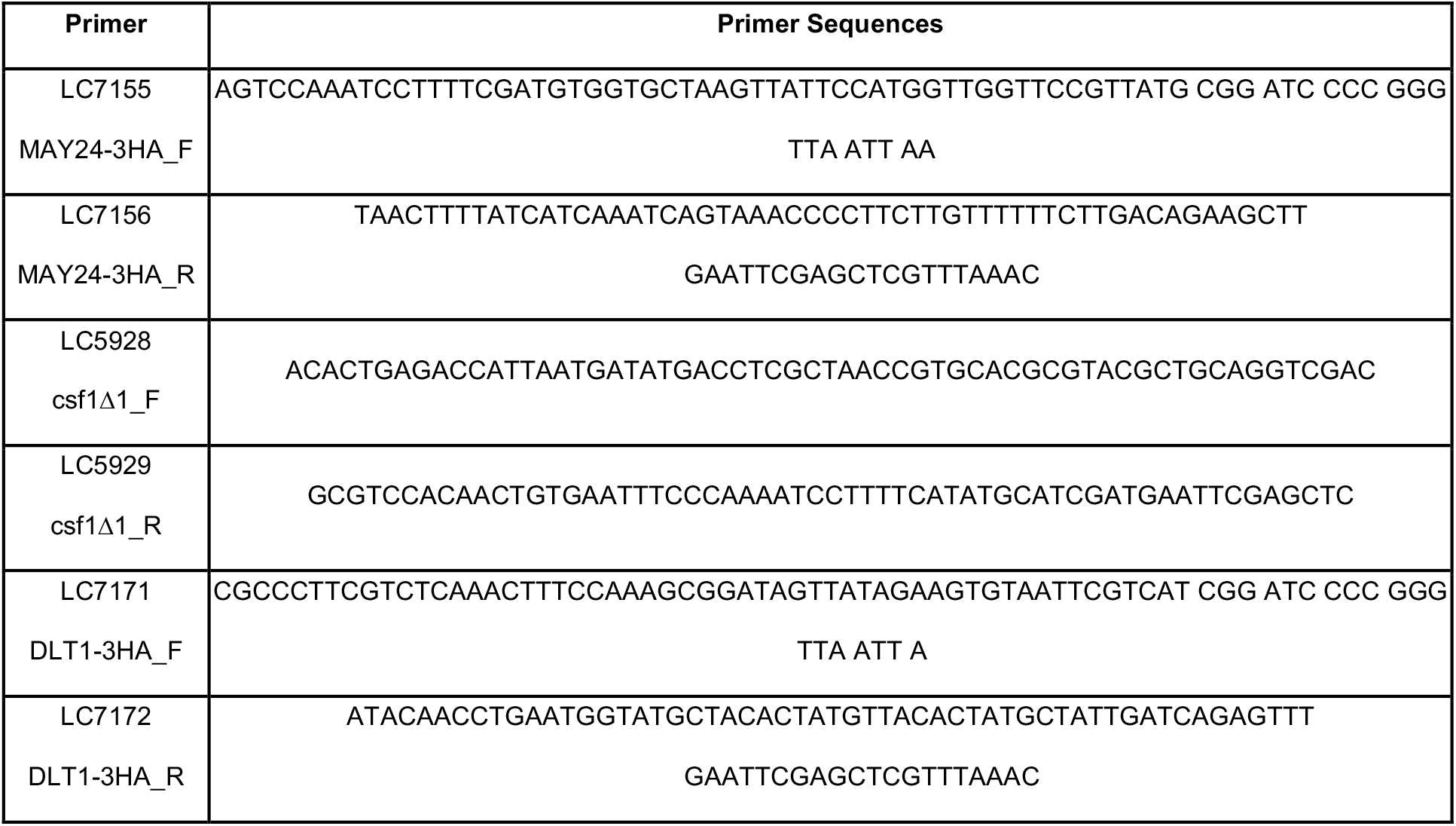
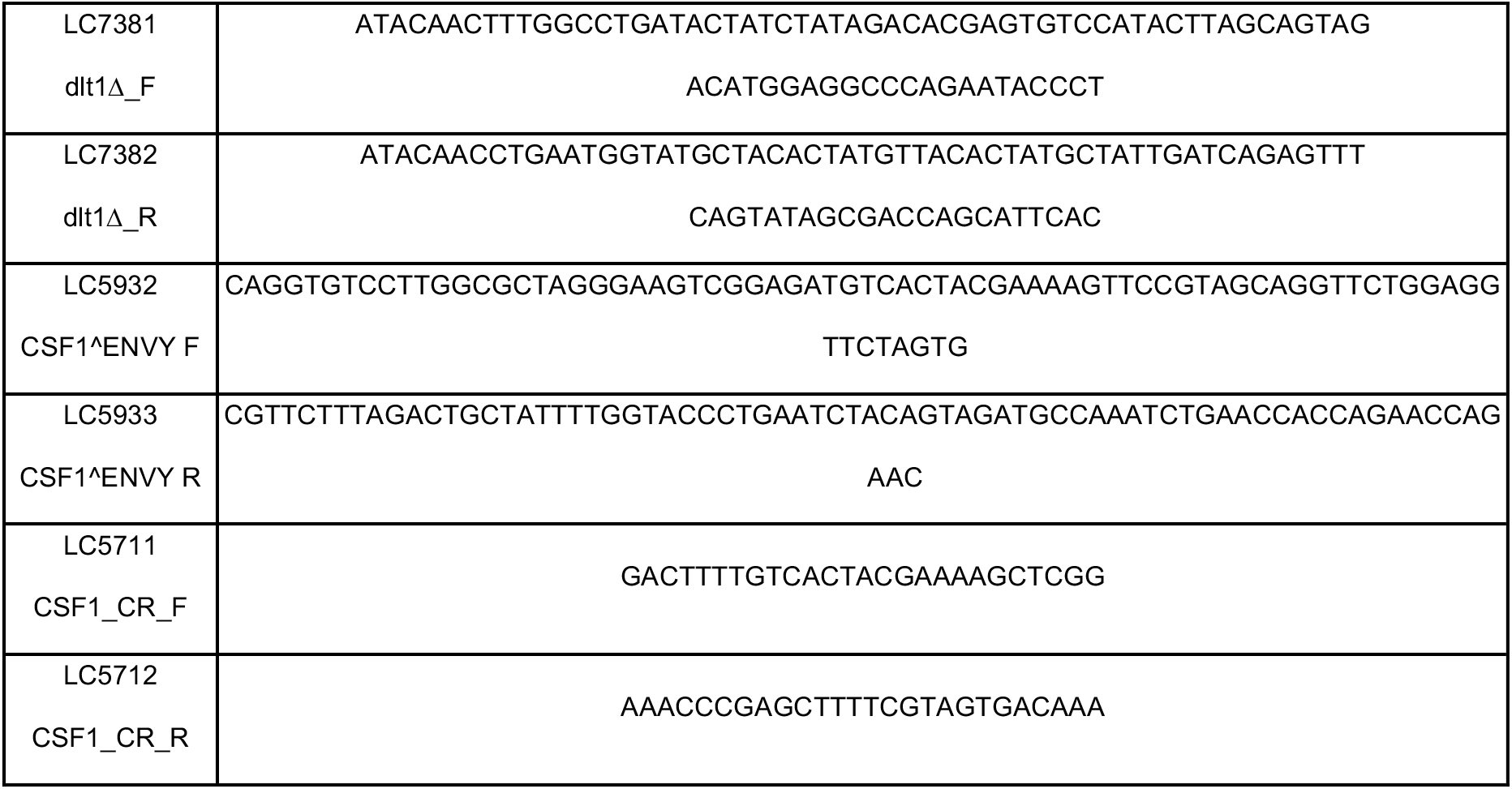
Yeast strains, plasmids and primers used in this study.

**Table S3.**
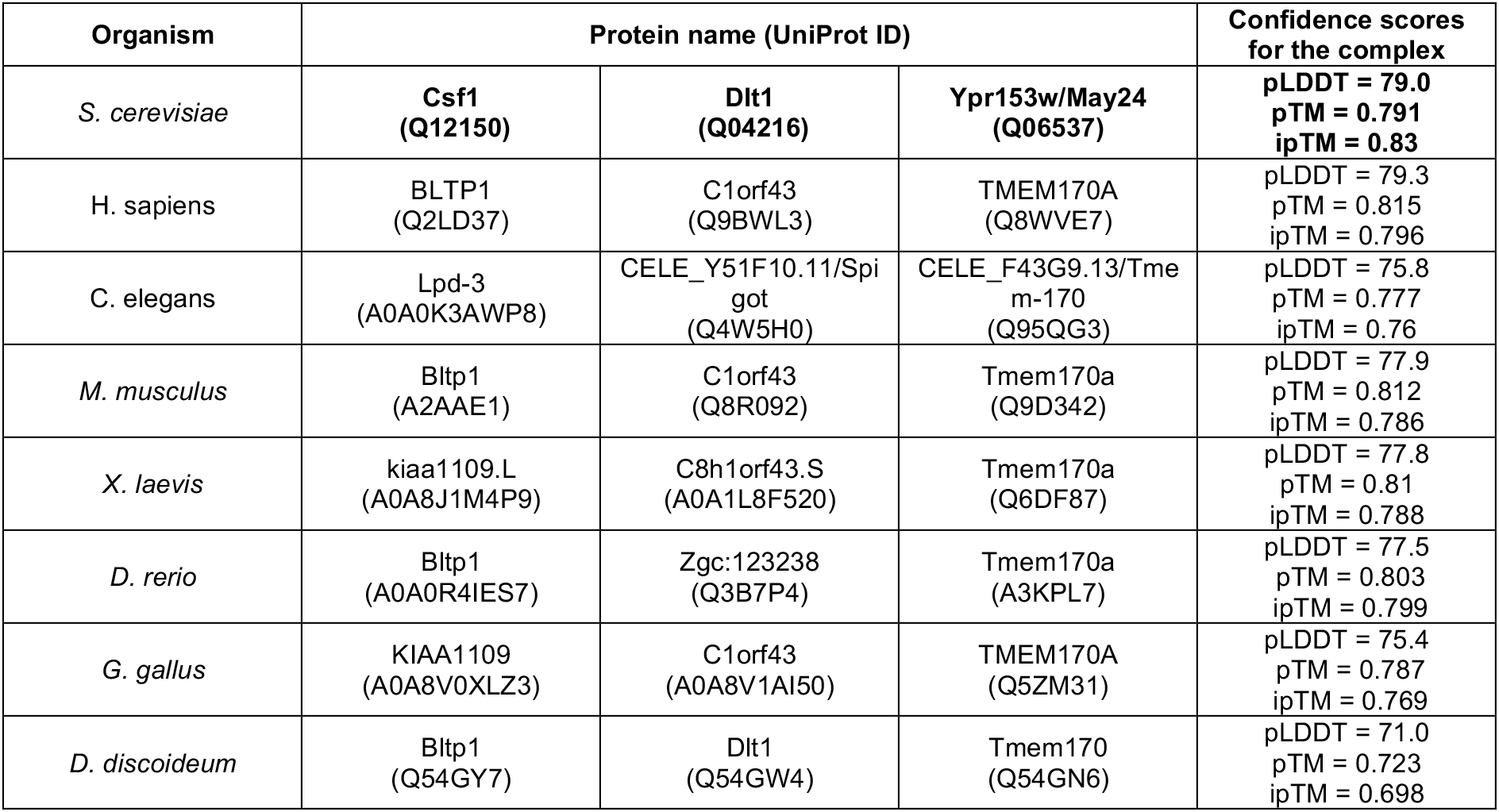
Homologues of fungal proteins Csf1, Dlt1 and May24 in different organisms. The UniProt identifier of each protein is shown in parentheses. The confidence scores correspond to the trimer predicted by AlphaFold.

| Organism | Protein name (UniProt ID) |  |  | Confidence scores for the complex |
| --- | --- | --- | --- | --- |
| *S. cerevisiae* | Csf1<br>(Q12150) | Dlt1<br>(Q04216) | Ypr153w/May24<br>(Q06537) | pLDDT = 79.0<br>pTM = 0.791<br>ipTM = 0.83 |
| *H. sapiens* | BLTP1<br>(Q2LD37) | C1orf43<br>(Q9BWL3) | TMEM170A<br>(Q8WVE7) | pLDDT = 79.3<br>pTM = 0.815<br>ipTM = 0.796 |
| *C. elegans* | Lpd-3<br>(A0A0K3AWP8) | CELE_Y51F10.11/Spi<br>got<br>(Q4W5H0) | CELE_F43G9.13/Tme<br>m-170<br>(Q95QG3) | pLDDT = 75.8<br>pTM = 0.777<br>ipTM = 0.76 |
| *M. musculus* | Bltp1<br>(A2AAE1) | C1orf43<br>(Q8R092) | Tmem170a<br>(Q9D342) | pLDDT = 77.9<br>pTM = 0.812<br>ipTM = 0.786 |
| *X. laevis* | kiaa1109.L<br>(A0A8J1M4P9) | C8h1orf43.S<br>(A0A1L8F520) | Tmem170a<br>(Q6DF87) | pLDDT = 77.8<br>pTM = 0.81<br>ipTM = 0.788 |
| *D. rerio* | Bltp1<br>(A0A0R4IES7) | Zgc:123238<br>(Q3B7P4) | Tmem170a<br>(A3KPL7) | pLDDT = 77.5<br>pTM = 0.803<br>ipTM = 0.799 |
| *G. gallus* | KIAA1109<br>(A0A8V0XLZ3) | C1orf43<br>(A0A8V1AI50) | TMEM170A<br>(Q5ZM31) | pLDDT = 75.4<br>pTM = 0.787<br>ipTM = 0.769 |
| *D. discoideum* | Bltp1<br>(Q54GY7) | Dlt1<br>(Q54GW4) | Tmem170<br>(Q54GN6) | pLDDT = 71.0<br>pTM = 0.723<br>ipTM = 0.698 |

## Supplementary figures

**Supplementary Figure S1.**
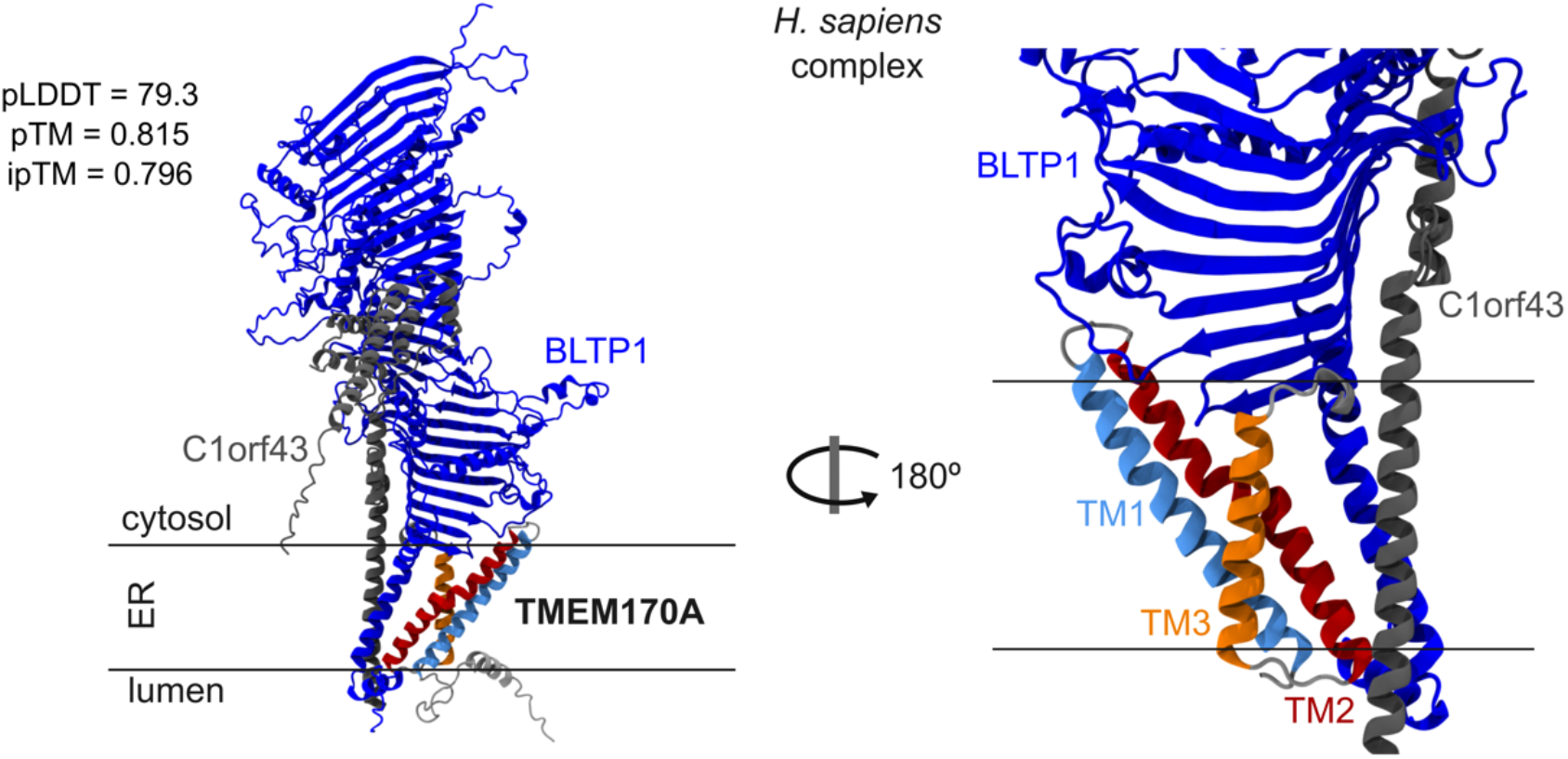
AF2 prediction of the heterotrimeric complex from *H. sapiens* formed by BLTP1 (blue, sequence from Met1 to Lys1200), C1orf43 (gray, sequence from Met1 to Pro225) and full length TMEM170A protein (the three TM helices are shown in light blue, red and orange, respectively.)

**Supplementary Figure S2.**
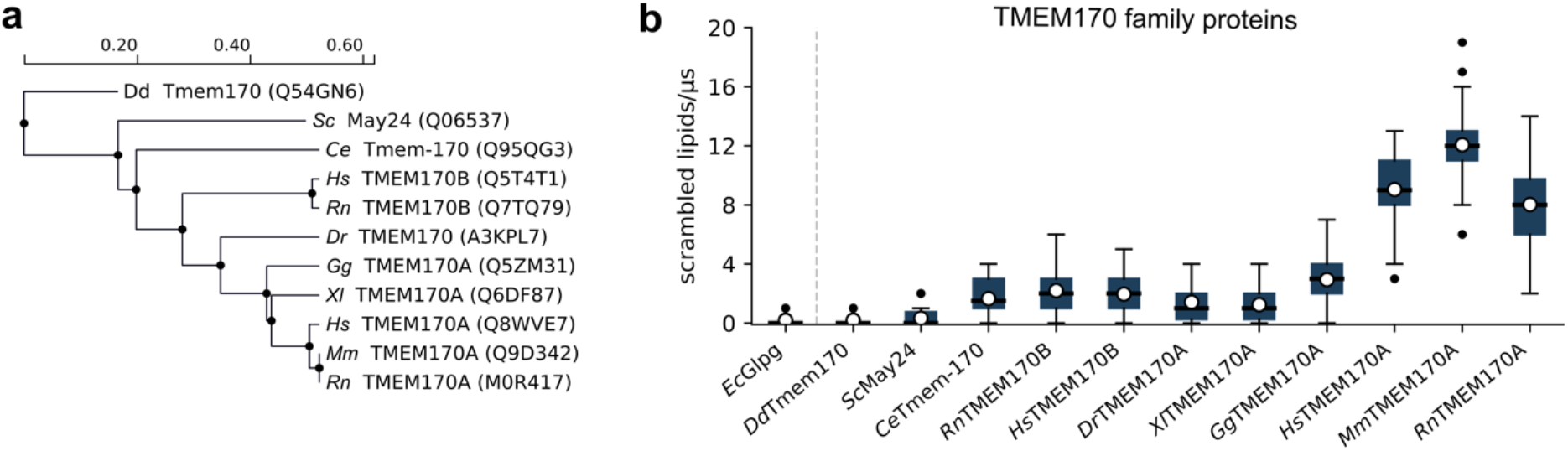
**a**. Phylogenetic tree depicting evolutionary relationships between the TMEM170 proteins used for this study. The scale shown on top of the tree represents the evolutive distance between sequences. The UniProt ID for each sequence is shown in parenthesis. **b**. *In silico* lipid scrambling for the selected TMEM170 proteins. A total of 46 data points were used to build each boxplot. The blue boxes represent the interquartile region. The solid black line inside each box represents the median of the data while the whiskers represent the data range. Averages are shown as white circles. Outliers are shown as black dots.

**Supplementary Figure S3.**
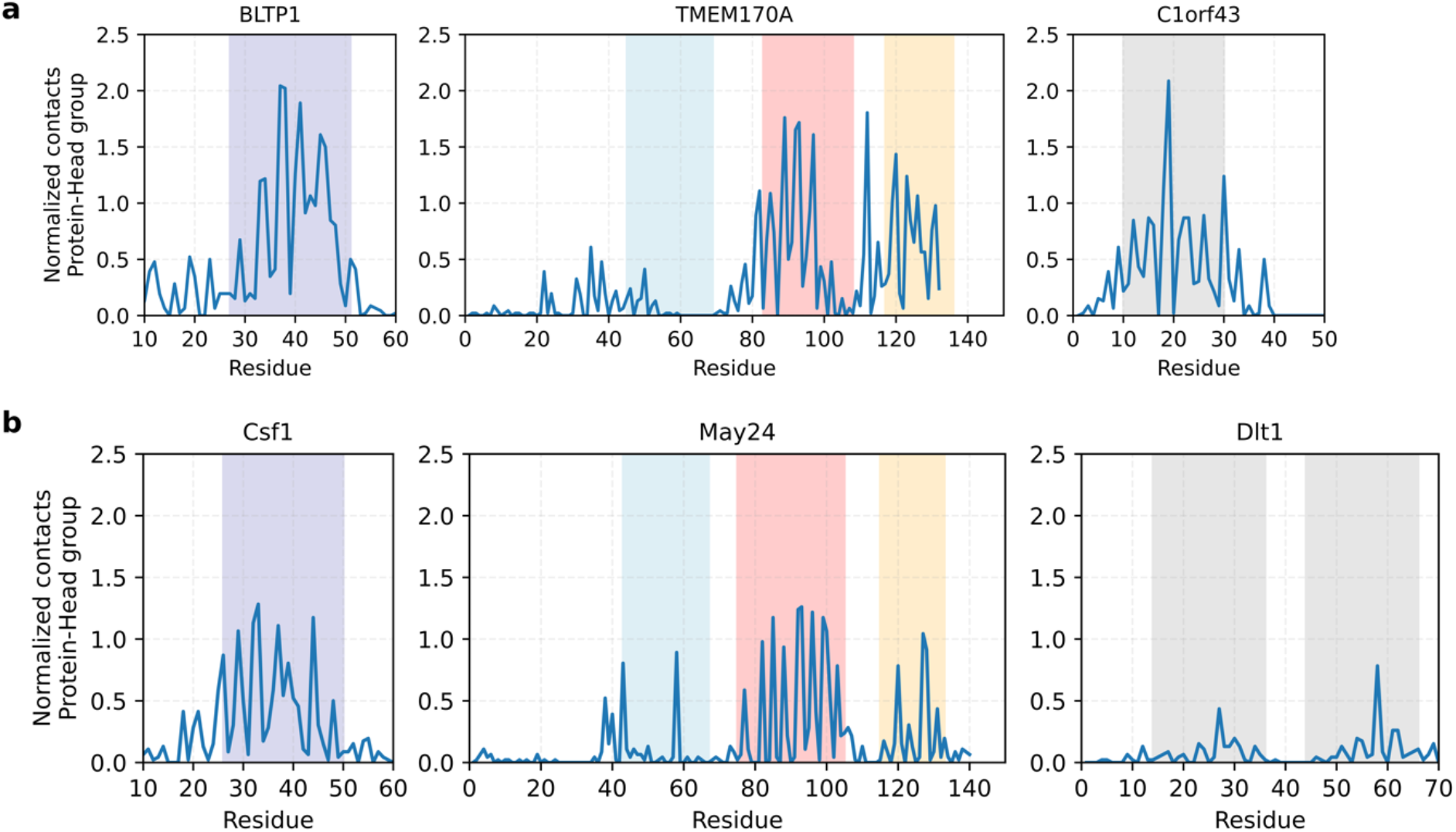
Interaction between the head groups of the scrambled lipids and each component from the heterotrimeric complex. **(a)** *H. sapiens* complex (Left panel: BLTP1; Center panel: TMEM170A ; Right panel: C1orf43). (**b**) *S. cerevisiae* complex (Left panel: Csf1; Center panel: May24 ; Right panel: Dlt1). The TM segments of BLTP1 and Csf1 are highlighted with blue shadows. The three TM segments of TMEM170A and May24 are highlighted as in panel **(a)** and Figure **1a**. The TM segments of C1orf43 and Dlt1 are highlighted with gray shadows.

## References

1. Gómez-Sánchez, R. et al. Atg9 establishes Atg2-dependent contact sites between the endoplasmic reticulum and phagophores. J. Cell Biol. 217, 2743–2763 (2018).

2. Matoba, K. et al. Atg9 is a lipid scramblase that mediates autophagosomal membrane expansion. Nat. Struct. Mol. Biol. 27, 1185–1193 (2020).

3. Ghanbarpour, A., Valverde, D. P., Melia, T. J. & Reinisch, K. M. A model for a partnership of lipid transfer proteins and scramblases in membrane expansion and organelle biogenesis. Proc. Natl. Acad. Sci. 118, e2101562118 (2021).

4. Reinisch, K. M.Chen, X.-W. & Melia, T. J. “VTT”-Domain Proteins VMP1 and TMEM41B Function in Lipid Homeostasis Globally and Locally as ER Scramblases. Contact 4, 251525642110244 (2021).

5. John Peter, A. T. et al. Vps13-Mcp1 interact at vacuole-mitochondria interfaces and bypass ER-mitochondria contact sites. J. Cell Biol. 216, 3219–3229 (2017).

6. Guillén-Samander, A. et al. A partnership between the lipid scramblase XK and the lipid transfer protein VPS13A at the plasma membrane. Proc. Natl. Acad. Sci. U. S. A. 119, e2205425119 (2022).

7. Park, J.-S., Hu, Y., Hollingsworth, N. M., Miltenberger-Miltenyi, G. & Neiman, A. M. Interaction between VPS13A and the XK scramblase is important for VPS13A function in humans. J. Cell Sci. 135, jcs260227 (2022).

8. Gao, J. et al. Any1 is a phospholipid scramblase involved in endosome biogenesis. J. Cell Biol. (2025) doi:10.1083/jcb.202410013.

9. Kurosaka, G. et al. A novel ER membrane protein Ehg1/May24 plays a critical role in maintaining multiple nutrient permeases in yeast under high-pressure perturbation. Sci. Rep. 9, 18341 (2019).

10. Abe, F. & Minegishi, H. Global Screening of Genes Essential for Growth in High-Pressure and Cold Environments: Searching for Basic Adaptive Strategies Using a Yeast Deletion Library. Genetics 178, 851–872 (2008).

11. Kang, Y., Lehmann, K.S., Long, H. et al. Structural basis of lipid transfer by a bridge-like lipid-transfer protein. Nature. (2025) doi: 10.1038/s41586-025-08918-y.

12. Christodoulou, A., Santarella-Mellwig, R., Santama, N. & Mattaj, I. W. Transmembrane protein TMEM170A is a newly discovered regulator of ER and nuclear envelope morphogenesis in human cells. J. Cell Sci. 129, 1552–1565 (2016).

13. Jamali, K., Käll, L., Zhang, R., Brown, A., Kimanius, D. & Scheres, S. H. W. Automated model building and protein identification in cryo-EM maps. Nature. 628, 450–457 (2024).

14. Li, D., Rocha-Roa, C., Schilling, M. A., Reinisch, K. M. & Vanni, S. Lipid scrambling is a general feature of protein insertases. Proc. Natl. Acad. Sci. U. S. A. 121, e2319476121 (2024).

15. Söding, J., Biegert, A. & Lupas, A. N. The HHpred interactive server for protein homology detection and structure prediction. Nucleic Acids Res. 33, W244–248 (2005).

16. UniProt Consortium. UniProt: the Universal Protein Knowledgebase in 2025. Nucleic Acids Res. 53, D609–D617 (2025).

17. Jumper, J. et al. Highly accurate protein structure prediction with AlphaFold. Nature 596, 583–589 (2021).

18. Evans, R. et al. Protein Complex Prediction with AlphaFold-Multimer. http://biorxiv.org/lookup/doi/10.1101/2021.10.04.463034 (2021) xdoi:10.1101/2021.10.04.463034.

19. Mirdita, M. et al. ColabFold: making protein folding accessible to all. Nat. Methods 19, 679– 682 (2022).

20. Lomize, M. A., Pogozheva, I. D., Joo, H., Mosberg, H. I. & Lomize, A. L. OPM database and PPM web server: resources for positioning of proteins in membranes. Nucleic Acids Res. 40, D370–D376 (2012).

21. Abraham, M. J. et al. GROMACS: High performance molecular simulations through multi-level parallelism from laptops to supercomputers. SoftwareX 1–2, 19–25 (2015).

22. Souza, P. C. T. et al. Martini 3: a general purpose force field for coarse-grained molecular dynamics. Nat. Methods 18, 382–388 (2021).

23. Kroon, P. et al. Martinize2 and Vermouth: Unified Framework for Topology Generation. Preprint at 10.7554/eLife.90627.2 (2024).

24. Wassenaar, T. A., Ingólfsson, H. I., Böckmann, R. A., Tieleman, D. P. & Marrink, S. J. Computational Lipidomics with insane: A Versatile Tool for Generating Custom Membranes for Molecular Simulations. J. Chem. Theory Comput. 11, 2144–2155 (2015).

25. Bussi, G., Donadio, D. & Parrinello, M. Canonical sampling through velocity rescaling. J. Chem. Phys. 126, 014101 (2007).

26. Parrinello, M. & Rahman, A. Polymorphic transitions in single crystals: A new molecular dynamics method. J. Appl. Phys. 52, 7182–7190 (1981).

27. Janke, C. et al. A versatile toolbox for PCR-based tagging of yeast genes: new fluorescent proteins, more markers and promoter substitution cassettes. Yeast Chichester Engl. 21, 947–962 (2004).

28. Longtine, M. S. et al. Additional modules for versatile and economical PCR-based gene deletion and modification in Saccharomyces cerevisiae. Yeast Chichester Engl. 14, 953–961 (1998).

29. Slubowski, C. J., Funk, A. D., Roesner, J. M., Paulissen, S. M., Huang, L. S. Plasmids for C-terminal tagging in Saccharomyces cerevisiae that contain improved GFP proteins, Envy and Ivy. Yeast. 32, 379–87 (2015).

